# Compressive stress drives adhesion-dependent unjamming transitions in breast cancer cell migration

**DOI:** 10.1101/2022.04.30.490153

**Authors:** Grace Cai, Anh Nguyen, Yashar Bashirzadeh, Shan-Shan Lin, Dapeng Bi, Allen P. Liu

**Affiliations:** Applied Physics Program, University of Michigan, Ann Arbor, MI, USA; Department of Physics, Northeastern University, Boston, MA, USA; Department of Mechanical Engineering, University of Michigan, Ann Arbor, MI, USA; Department of Biomedical Engineering, University of Michigan, Ann Arbor, MI, USA; Department of Biophysics, University of Michigan, Ann Arbor, MI, USA; Cellular and Molecular Biology Program, University of Michigan, Ann Arbor, MI, USA

## Abstract

Cellular unjamming is the collective fluidization of cell motion and has been linked to many biological processes, including development, wound repair, and tumor growth. In tumor growth, the uncontrolled proliferation of cancer cells in a confined space generates mechanical compressive stress. However, because multiple cellular and molecular mechanisms may be operating simultaneously, the role of compressive stress in unjamming transitions during cancer progression remains unknown. Here we investigate which mechanism dominates in a dense, mechanically stressed monolayer. We find that long-term mechanical compression triggers cell arrest in benign epithelial cells and enhances cancer cell migration in transitions correlated with cell shape, leading us to examine the contributions of cell-cell adhesion and substrate traction in shape-dependent unjamming transitions. We show that cadherin-mediated cell-cell adhesion regulates differential cellular responses to compressive stress and predominantly controls unjamming transitions in dense monolayers. Importantly, compressive stress does not induce the epithelial—mesenchymal transition in unjammed cells. Using traction force microscopy, traction forces are attenuated in compressed cells in the bulk of the monolayer regardless of cell type and motility. Intercellular adhesion thus is the dominant regulator of compression-induced unjamming transitions and may impact collective cell motion in tumor development and breast cancer progression.

## Introduction

Cellular jamming impacts many fundamental biological and disease processes including embryogenesis (Alert and Trepat, 2020), tissue repair (Nnetu *et al*., 2012; Ajeti *et al*., 2019), and tumor growth (Wang *et al*., 2020; Gensbittel *et al*., 2021). A jamming transition is a transition from a solid-like state to a fluid-like state in which cellular rearrangements are diminished. Jamming transitions during embryogenesis are typically governed by cell density (Blauth *et al*., 2021). As a monolayer ages, cells proliferate, slow down, and become jammed as a dense cell layer. Jammed cells are observed to be confined in “cages” of the size of a single cell by their neighbors (Garcia *et al*., 2015). The friction between cells increases, leading to reduced collective and individual motion. A dense monolayer in a solid-like state can quickly revert to a flowing state when a wound is inflicted (Chepizhko *et al*., 2018). During wound repair, the ability for cells to rearrange is essential for closing gaps in epithelial tissues and may be regulated by jamming transitions. Although jamming plays a crucial role in many biological events, the main parameters of cellular jamming remain poorly understood.

Recent research suggests cellular unjamming is involved in tumor growth and cancer progression (Haeger *et al*., 2014; Staneva *et al*., 2019; Grosser *et al*., 2021). During tumor growth, cancer cells proliferate in a dense and confined environment, subjecting tumor cells to solid compressive stress (Northcott *et al*., 2018). Solid stresses affect tumor pathophysiology by directly compressing cancer and stromal cells and indirectly deforming blood and lymphatic vessels (Jain, Martin and Stylianopoulos, 2014). For tumors to grow and proliferate, cancer cells must be able to move and divide. Cells in parts of the tumor can fluidize and migrate collectively in an unjamming transition^4^. Here we investigate the idea that unjamming transitions in cancer cells are driven by compressive stress. As solid stress increases within a tumor, microenvironmental factors may prime cells toward invasive phenotypes, giving rise to cellular rearrangements and enhanced migration^6,7^.

Cellular rearrangements cease when the cell shape index approaches a critical value (Park *et al*., 2016). Based on the vertex model, which defines a shape index 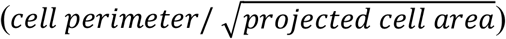 of 3.81 as the jamming threshold (Park *et al*., 2015), structural rearrangement requires cell shape changes. In this way, cells can overcome the jamming constraints of density by adapting their shapes. Densely packed cells can still move if they elongate (i.e., increase shape index above 3.81). As the cell shape index decreases and approaches the critical value, the tissue becomes jammed. Thus, the jamming transition can be controlled by the preferred cell shape.

It is known that cell-cell and cell-substrate adhesion forces act together to generate a preferred cell shape (Farhadifar *et al*., 2007; Bi *et al*., 2016; Trepat and Sahai, 2018). Adhesions are major sites of force transmission in cells and generally strengthen as cells approach jamming (Garcia *et al*., 2015). Cell-cell adhesion is mediated by cadherins that are anchored to the cytoskeleton (Charras and Yap, 2018), whereas integrin-dependent cell-substrate adhesion is governed by focal adhesions that generate internal cytoskeletal tension (Kanchanawong *et al*., 2010). In nonconfluent tissues, decreasing cell-cell adhesion reduces cell crowding and cell-cell contacts, increasing the fluidity of the tissue (Lawson-Keister and Manning, 2021). In confluent tissues, the role of cell-cell adhesion tends to be cell-type specific and dependent on invasive potential (Tse *et al*., 2012). Strong cell-substrate adhesion combined with high traction stresses are shown to contribute to unjamming in confluent systems (Malinverno *et al*., 2017). Relatively small changes in cell-cell and cell-substrate adhesion can have profound effects on tissue rheology and can be used to regulate cell arrest (Park *et al*., 2016; Rens and Merks, 2020). How cell-cell and cell-substrate adhesion manipulate jamming transitions in a dense, mechanically stressed monolayer remains unclear.

Here we characterize the role adhesion complexes play in regulating differential cellular responses to mechanical compression and in driving cellular unjamming transitions in cancer progression. As normal breast epithelial cells (MCF10A) and metastatic breast cancer cells (4T1) are subjected to compressive stress, MCF10A cells fluidize while 4T1 cells behave solid-like. In the process, 4T1s elongate and develop strong cell-cell adhesions. In contrast, following compression, E-cadherin is disrupted at the cell-cell contacts of MCF10As, which become stationary and acquire a more compact cell shape. As mesenchymal markers are not upregulated in compressed 4T1s, the transition is distinct from the epithelial-to-mesenchymal transition (EMT). We show that E-cadherin-mediated cell-cell adhesion is the dominant regulator of the observed differential cellular responses to mechanical compression and predominantly controls cellular jamming and unjamming transitions.

## Results

### Long-term compressive stress drives cellular jamming and unjamming transitions

We first asked whether the effects of long-term mechanical compression on collective cell migration depended on invasive potential. To address this, we conducted wound healing assays of non-tumorigenic (MCF10A) and cancer (4T1) breast epithelial cells subjected to different levels of compressive stress. The time evolution of the wound margin was monitored over 16 hours (**Fig. 1A**). Confluent MCF10A and 4T1 monolayers were scratched to create a uniform wound, inducing migration, and normal compressive force was applied to the cells. This in vitro model has been used previously by us and others and mimics the solid compressive stress experienced by cells during tumor development (Tse *et al*., 2012; Luo *et al*., 2022).

**Figure 1.**
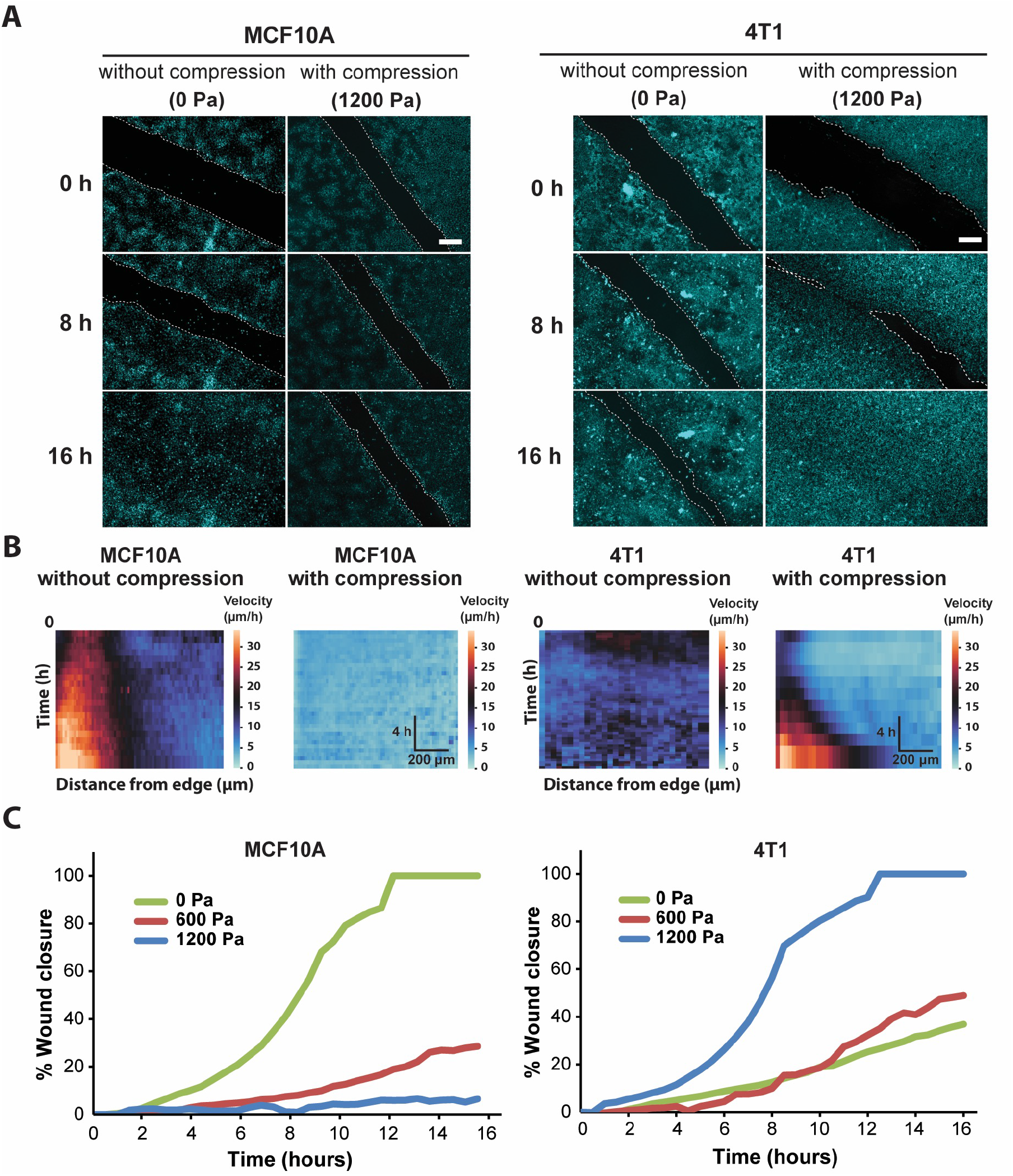
Compressive stress inhibits migration in MCF10A cells and enhances migration in 4T1 cells. **(A)** Representative fluorescence images of MCF10A and 4T1 wound area at the indicated time points post-wound with and without compression. Cell nuclei are labeled with Hoechst 33342. Lines are scratched on each well using a p-200 pipette tip and relative cell migration is captured by fluorescence microscopy at 30 min time intervals for 16 h post-wound. Cell edges used to calculate wound area are outlined by white dashed lines. Scale bars, 50 µm. **(B)** Heat maps show spatiotemporal evolution of the velocity for control and compressed cells. **(C)** Quantification of wound area (between white dashed cell edges) for each cell type and compressive pressure. Mean wound area at each time point is plotted from 3 independent replicates as a representative trace.

During wound healing, the entire cell sheet of control MCF10A cells moved collectively at a constant velocity to close the wound (Supplementary Movie 1). The epithelial sheet was in a seemingly motile but locally jammed state as collective motion was high, however only cells near the wound edge exhibited high motility, indicating that cells in the bulk of the monolayer were caged by their neighbors (**Fig. 1B**). In contrast, cell velocity in control 4T1 monolayer was low, leading to minimal collective migration and failure to close the wound (**Fig. 1B** and Supplementary Movie 2). When MCF10A monolayer was exposed to compressive stresses of 600 and 1,200 Pa, the cells showed very low migratory ability (**Fig. 1B** and Supplementary Movie 3). However, when 4T1 monolayer was subjected to 1,200 Pa of compressive stress, the cells underwent highly collective, fluid-like migration and abruptly closed the wound after approximately 8 hours (**Fig. 1A** and Supplementary Movie 4). By applying different levels of mechanical compression and tracking the wound margin over time, we demonstrated that compressive stress attenuated cell motility in MCF10A cells, which quickly entered a jammed state. 4T1 cells reacted actively to compressive stress by transitioning to a fluid-like state (**Fig. 1C**). Enhanced collective migration of 4T1 cells was positively correlated with the level of external stress applied.

### Unjamming is linked to changes in cell shape and nuclear shape

We next investigated cell shape as a marker for tissue fluidity in dense tissues. Previously published work used the vertex model to define a critical shape index of 3.81 as the jamming threshold (Bi *et al*., 2016). Studies using human bronchial epithelial cells show that regardless of the magnitude of intracellular stress fluctuations, cellular rearrangements cease when the cell shape index approaches the jamming threshold (Park *et al*., 2015).

We examined cell shape as a parameter of compression-induced jamming and unjamming transitions in a dense monolayer. When compressive stress was applied to the MCF10A monolayer, cell shape index was reduced, approaching the critical value of 3.81, also known as the jamming threshold (**Fig. 2A and 2B**). MCF10A cells reacted to compressive stress by becoming more compact (lowered shape index). When 4T1 cells were subjected to compression, cellular and nuclear shapes became more elongated (higher shape index), and more variable compared to control cells (**Fig. 2B**). Although cell shape index depends on elongation and tortuosity, cell aspect ratio (AR) emphasizes elongation and deemphasizes tortuosity (Mitchel *et al*., 2020). Thus, cellular elongation can be better captured by plotting the mean of AR vs. the standard deviation (s.d.) of AR. We found this to have a positive linear relationship (**Fig. 2C**), in agreement with what has been shown previously (Atia *et al*., 2018). As cell AR increased with compressive stress, indicative of unjamming, its variability from cell to cell increased as well. Compressed cells tended to have higher values of 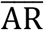 and s.d. of AR. Increased cellular elongation and shape variability suggest more disordered cell packing and fluid-like behavior and may be an indicator for increased metastatic potential. These results, together with the earlier wound healing data, show that jamming and unjamming transitions in MCF10A and 4T1 cells depend on cell shape.

**Figure 2.**
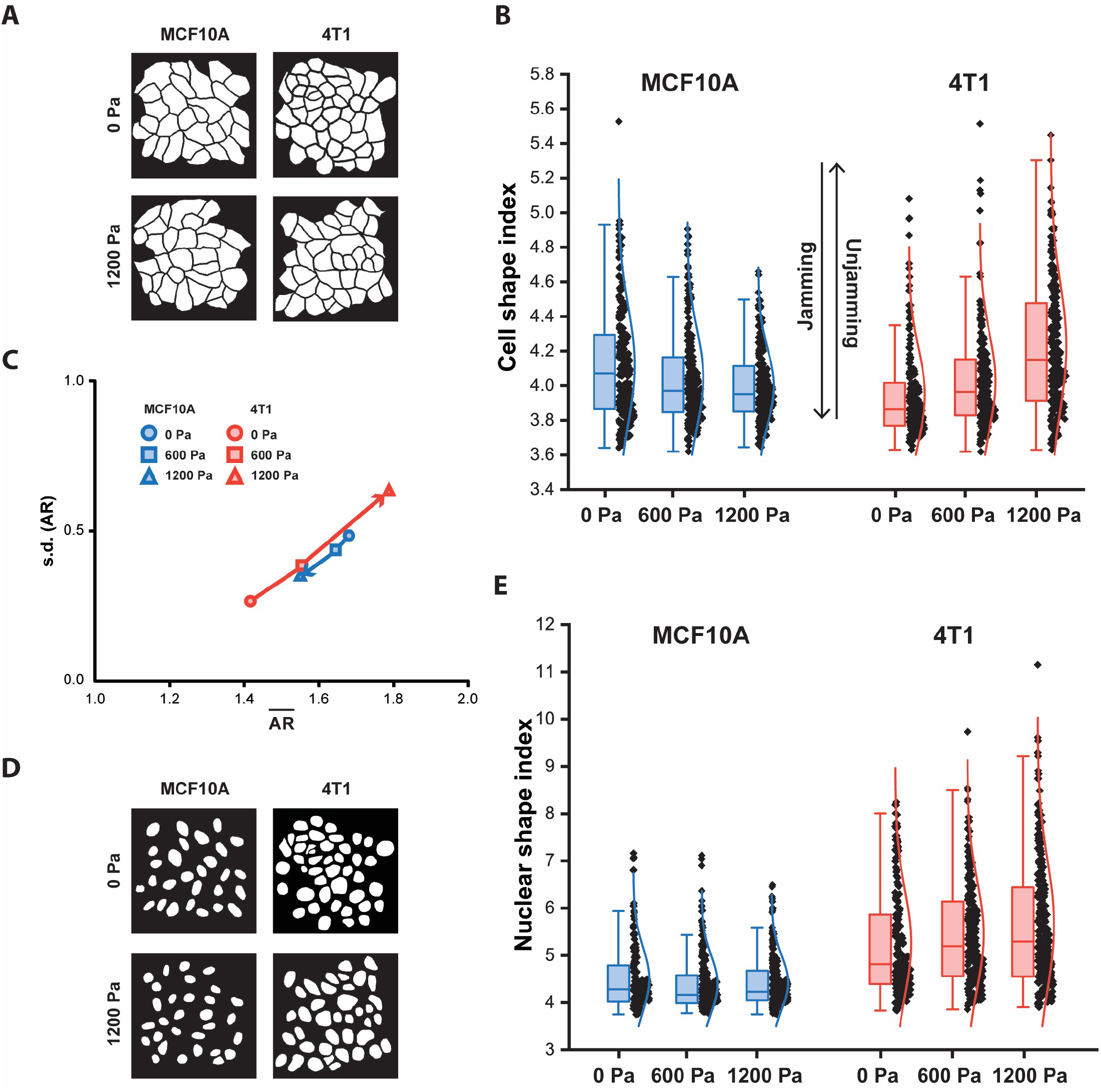
With compression, MCF10A cells and nuclei become more compact, whereas 4T1 cells and nuclei become elongated. **(A)** Representative binary images outlining fixed MCF10A and 4T1 cells labeled with E-cadherin with and without compression. **(B)** Boxplot of cell shape index for MCF10A and 4T1 cells subjected to 0, 600 and 1,200 Pa mechanical compression for 12 h. **(C)** Cell aspect ratio (AR), which emphasizes elongation, of control and compressed MCF10A and 4T1 cells is plotted as the mean of AR 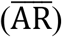 vs. the standard deviation (s.d.) of AR for each cell type and compressive pressure. **(D)** Representative binary images outlining fixed MCF10A and 4T1 cell nuclei labeled with DAPI with and without compression. **(E)** Boxplot of nuclear shape index for control and compressed MCF10A and 4T1 cells. Boxplots of cell and nuclear shape indices show median and quartiles for 3 independent replicates. Number of MCF10A cells and nuclei analyzed for compressive pressures: 0 Pa (n = 234), 600 Pa (n = 229), 1200 Pa (n = 222). Number of 4T1 cells and nuclei analyzed for compressive pressures: 0 Pa (n = 205), 600 Pa (n = 231), 1200 Pa (n = 224).

Along with cell shape, nuclear shape has recently also been linked to tissue fluidity (Grosser *et al*., 2021) and is a critical marker for tumor aggressiveness in clinical cancer grading (Denais and Lammerding, 2014). Cancer cell nuclei are generally larger and softer than non-malignant cell nuclei (Fischer, Hayn and Mierke, 2020; Rianna, Radmacher and Kumar, 2020; Gensbittel *et al*., 2021). Studies of multiple cancer cell types, including breast cancer cells, have found that the cells and their nuclei become significantly softer upon extravasation (Roberts *et al*., 2021). Since nucleus deformability is known to play a central role in cell motility in dense environments (Friedl, Wolf and Lammerding, 2011), we next asked whether changes in nuclear shape were correlated with changes in cell shape in cells subjected to mechanical compression. As expected, compressive stress reduced MCF10A nuclear shape index. As MCF10A cell shape index decreased with compressive stress, cell nuclei also became more compact (**Fig. 2D**). Elongated cell shapes in compressed 4T1 cells corresponded to increased nuclear shape index and high variance in nuclear shape (**Fig. 2E**), which has been associated with more aggressive tumors (Grosser *et al*., 2021). Our results show that cellular and nuclear shape indices increase with compressive stress in unjamming transitions and are important indicators of cell motility and tissue fluidity. More importantly, mechanical compression resulted in more elongated cell shapes in metastatic 4T1 cells, which became unjammed, but not in non-tumorigenic, jammed MCF10A cells.

### Compression-induced unjamming transition is distinct from EMT

To further elucidate how compressive stress impacts cell-cell organization, we next examined the factors which shape cells in a dense monolayer and support cell migration. We first investigated the effect of cell-cell adhesion after long-term compression. Immunofluorescence staining revealed compressive stress disrupted E-cadherin localization at the cell-cell contacts of MCF10A cells (**Fig. 3A and 3B**). Surprisingly, 4T1 cells gained strong cell-cell contacts evidenced by increased recruitment of E-cadherin to cell junctions when compressed (**Fig. 3C and 3D**). We hypothesized that the lack of collective migration exhibited by 4T1 cells in the absence of applied stresses can be attributed to weak cell-cell adhesion. High adhesion forces in compressed 4T1s encouraged individual cells to contribute equally to collective movement, and the cell layer became fluidized. This result supports that adhesion strength is increased in a more unjammed monolayer.

**Figure 3.**
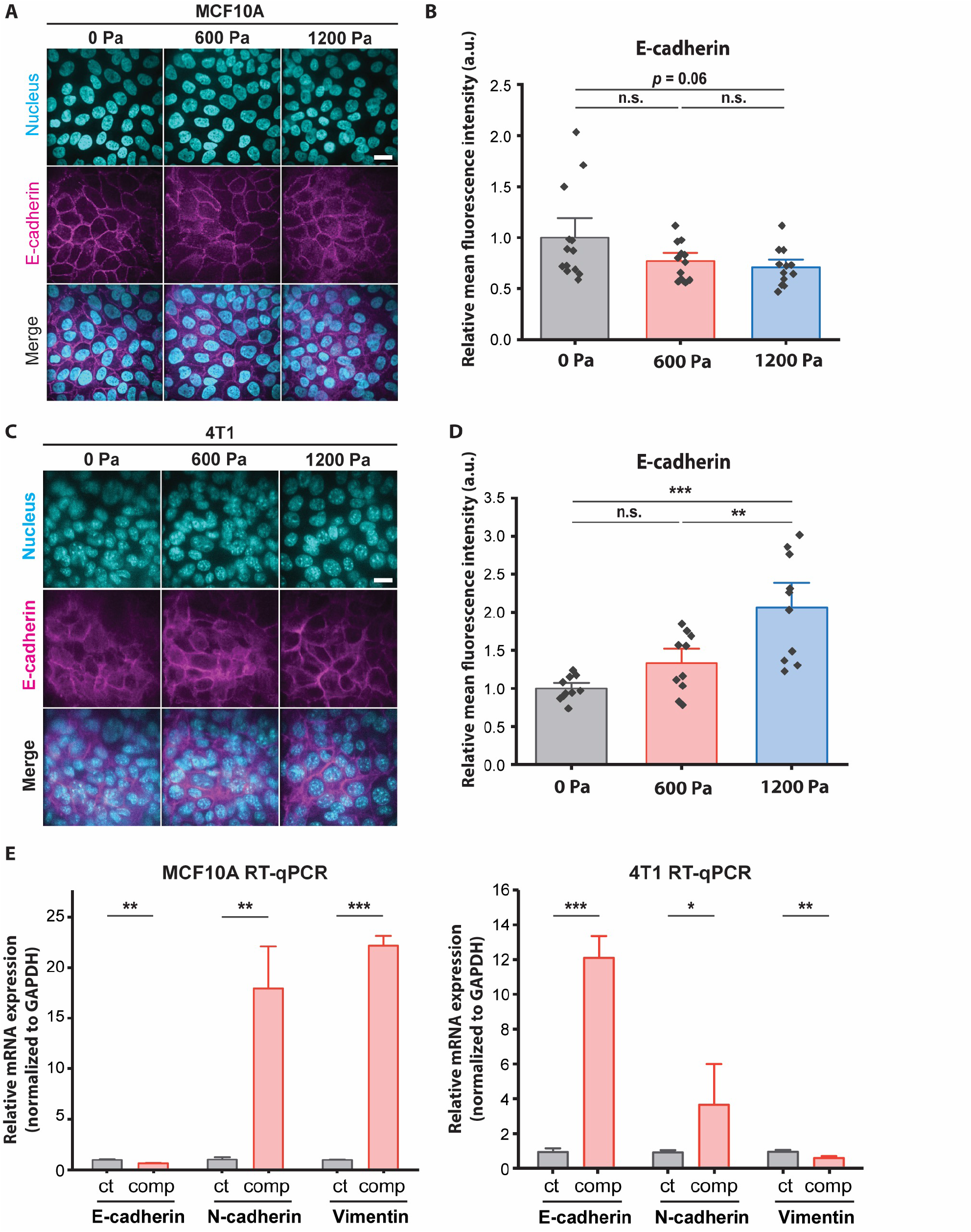
E-cadherin is upregulated in unjammed 4T1 cells under mechanical compression. **(A)** Representative immunofluorescence images of MCF10A cells labeled with DAPI and an E-cadherin antibody. Cell monolayers are subjected to specified compressive pressures for 12 h. Scale bar, 20 µm. **(B)** Quantification of relative E-cadherin fluorescence at MCF10A cell-cell contacts. Mean fluorescence intensity at the cell membrane ± S.D. is plotted from 3 independent replicates (n = 12-13). **(C)** Representative microscopy images of 4T1 cells labeled with DAPI and an E-cadherin antibody in the same experimental conditions as in (A). Scale bar, 20 µm. **(D)** Quantification of relative E-cadherin fluorescence at 4T1 cell-cell contacts. Mean fluorescence intensity at the cell membrane ± S.D. is plotted from 3 independent experiments (n = 10). **(E)** qPCR analysis of E-cadherin, N-cadherin, and vimentin mRNA levels with and without compression (1,200 Pa). Transcript levels are calculated using the ΔΔC_t_ method normalized to GAPDH. Mean mRNA level ± S.D. is plotted from 3 independent experiments with duplicates per experiment.

To identify a possible mesenchymal molecular signature in compressed cells that may attribute increased migratory ability to EMT^13–15^, we conducted qPCR assays to quantify mRNA levels of epithelial marker E-cadherin and mesenchymal markers N-cadherin and vimentin. Compressive stress significantly downregulated E-cadherin in MCF10A cells (**Fig. 3E and 3F**), while vimentin was upregulated. Surprisingly, E-cadherin was upregulated in compressed 4T1 cells by 11-fold, while vimentin was downregulated **(Fig. 3F)** – these are features associated with mesenchymal— epithelial transition (MET), as opposed to EMT. N-cadherin was upregulated by compression in both cell types, although mRNA level of MCF10A cells increased more substantially. Altogether, we find that enhanced cell motility and a higher cell shape index are associated with increased levels of cell-cell adhesion **(Fig. 2 and 3)**. The upregulation of E-cadherin and downregulation of vimentin in unjammed 4T1 cells suggest that the factors governing jamming transitions in this system is distinct from transitions between epithelial and mesenchymal cells.

### Cadherin-mediated cell-cell adhesion is required for unjamming

To probe the role of E-cadherin further, we attenuated cell-cell adhesions by knocking down E-cadherin (encoded by CDH1 gene) in 4T1 cells and tested the effect of knockdown on compression-induced unjamming (**Fig. 4A and 4B**). Upregulated E-cadherin in unjammed 4T1 cells suggests a dominant role for cell-cell adhesion in migrating monolayers under mechanical compression, and knockdown of E-cadherin is known to switch the migration mode of cells from collective to single-cell migration^16^. Although E-cadherin expression was still upregulated by compression after knockdown, upregulation was substantially reduced in 4T1 E-cadherin knockdown (E-cad KD) cells relative to wild-type (WT) cells (**Fig. 4C**). Applied stresses of 600 and 1,200 Pa reduced the motility of E-cad KD cells (**Fig. 4D and 4E**). In the absence of mechanical compression, both WT and E-cad KD cells achieved wound closure of ∼30% over 15 hours. Compression promoted wound closure for WT cells (∼100% for 1,200 Pa) while inhibiting wound closure in E-cad KD cells (∼6% for 1,200 Pa) (**Fig. 4F and Fig. 1C**). Our results indicate that E-cad KD cells existed in a jammed state when compressed, similar to what we observed in MCF10As.

**Figure 4.**
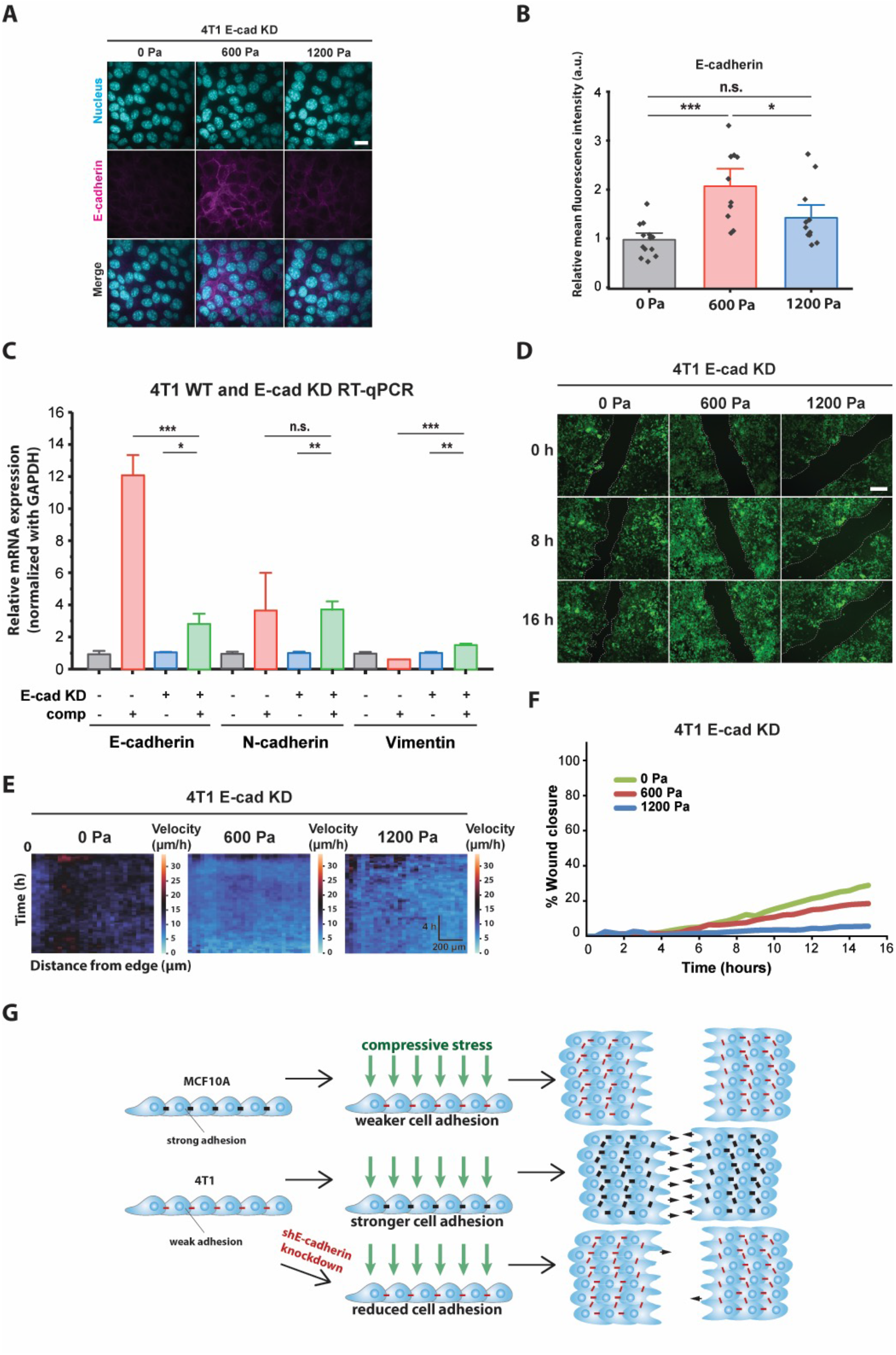
E-cadherin knockdown inhibited compression-induced upregulation of E-cadherin in 4T1 cells, triggering jamming. **(A)** Representative microscopy images of 4T1 E-cad KD cells labeled with DAPI and an E-cadherin antibody. E-cad KD is induced in 4T1 shE-cadherin cells by adding 200 μM IPTG 72 h prior to experiments. Cell monolayers are exposed to specified stresses for 12 h. Scale bar, 20 µm. **(B)** Quantification of relative E-cadherin fluorescence at the cell-cell contacts of 4T1 E-cad KD cells. Mean fluorescence intensity at the cell membrane ± S.D. is plotted from 3 independent replicates (n = 10) **(C)** qPCR analysis of E-cadherin, N-cadherin, and vimentin mRNA levels in 4T1 WT and 4T1 E-cad KD cells. Transcript levels >Luocalculated using the ΔΔC_t_ method normalized to GAPDH. Mean mRNA level ± S.D. is plotted from 3 independent experiments with duplicates per experiment. **(D)** Representative images of 4T1 E-cad KD wound area at the indicated time points post-wound. 4T1 shE-cadherin cells express mNeonGreen. Cell edges used to calculate wound area are outlined by white dashed lines. Scale bar, 50 µm. **(E)** Heat maps of spatiotemporal evolution of the velocity for 4T1 E-cad KD cells under different levels of mechanical compression. **(F)** Quantification of wound area (between white dashed cell edges) for each condition. Mean wound area at each time point is plotted from 3 independent replicates as a representative trace. **(G)** Summary depicting the effect of compressive stress on cell migration in MCF10A WT, 4T1 WT, and 4T1 E-cad KD cells. Strong cell-cell contacts are denoted by black dashes. Red dashes indicate weak cell-cell adhesion. Number of small black arrows (right) represent relative cell velocity in closing the wound.

To assess the generality of our finding, we probed the effect of mechanical compression on the cell motility of nonmetastatic mouse breast cancer cell line 67NR, which is derived from the same primary breast cancer as 4T1 and expresses N-cadherin and vimentin, but not E-cadherin (Dykxhoorn *et al*., 2009). Although 67NR cells have been shown to exhibit increased cell motility attributed to higher cell-substrate adhesion under mechanical compression (Tse *et al*., 2012), we did not observe this unjamming behavior in compressed 67NR cells (**Supplementary Fig. 1A**), suggesting that expression and localization of E-cadherin is required for compression-induced unjamming. Consistent with our findings for MCF10A, compressive stress did not increase E-cadherin levels in 67NR cells (**Supplementary Fig. 1B**). Based on our findings, schematically summarized in **Fig. 4G**, that compressive stress significantly hindered the coordinated migration of 4T1 E-cad KD cells, and that upregulation of E-cadherin is required for compression-induced unjamming, we conclude that E-cadherin-dependent cell-cell adhesion is a key regulator and effector upon compression.

### Compressive stress reduces traction forces of follower cells in jamming—unjamming transitions

We have shown thus far that increased cell-cell adhesion promotes the unjamming of densely packed cancer cells. Since high traction stresses have been shown to contribute to the unjamming of a confluent monolayer (Malinverno *et al*., 2017), we next explored the role of substrate traction as a potential parameter working together with cell-cell adhesion to promote unjamming in tissues. Vinculin is a cytoskeletal protein responsible for regulating integrin-mediated cell adhesion and is found in focal adhesions as well as adherens junctions (Kanchanawong *et al*., 2010). We found that compressive stress enhanced the enrichment of vinculin at adherens junctions in 4T1 cells seeded on micropatterned substrates (**Fig. 5A and 5B**). This is consistent with increased localization of E-cadherin (**Supplementary Fig. 2**) since vinculin also binds E-cadherin and is needed to form cell-cell contacts. Since an increase in vinculin intensity can result from higher cell-cell and/or cell-matrix adhesion, we turned to traction force microscopy (TFM)^19^ to disentangle the individual contributions of cell-cell adhesion and substrate traction.

**Figure 5.**
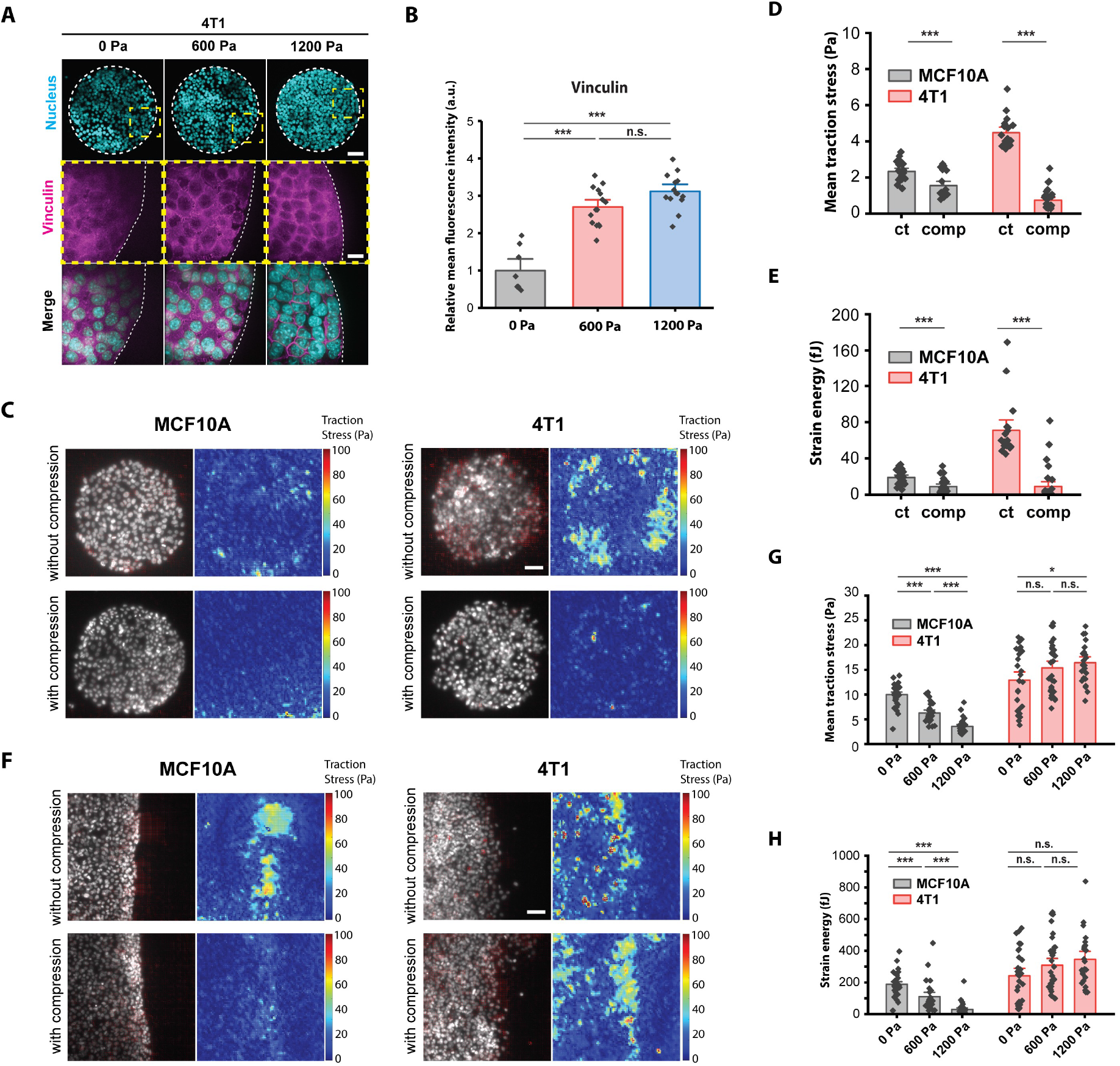
Compressive stress reduces traction forces of follower cells in MCF10A and 4T1 cell islands. **(A)** Representative microscopy images of 4T1 cell islands labeled with DAPI and a vinculin antibody. Micropatterned cell islands are exposed to specified compressive pressures for 12 h. Scale bars, 80 µm (top) and 20 µm (bottom). **(B)** Quantification of relative vinculin intensity. Mean fluorescence ± S.D. is plotted from 3 independent replicates (n = 8-15). **(C)** Traction stress vector field and traction stress magnitude of micropatterned cell islands with and without compression. Cell nuclei are labeled with Hoechst 33342. Compressed cell islands are subjected to 1,200 Pa applied stress for 3 h. Scale bar, 80 µm. **(D)** Mean traction stresses exerted by cells with and without compression on micropatterned islands. **(E)** Mean strain energy of cells with and without compression on micropatterned islands. Number of MCF10A cell islands analyzed: control (n = 21), compressed (n = 21). Number of 4T1 cell islands analyzed: control (n = 18) and compressed (n = 30). **(F)** Traction stress vector field and traction stress magnitude of a wounded edge with and without compression. Cell nuclei are labeled with Hoechst 33342. Compressed cells are subjected to 1,200 Pa applied stress for 3 h. Scale bar, 80 µm. **(G)** Mean traction stresses exerted by cells with and without compression at a wounded edge. **(H)** Mean strain energy of cells with and without compression at a wounded edge. Number of wounded MCF10A cell edges analyzed: 0 Pa (n = 37), 600 Pa (n = 26), 1,200 Pa (n = 32). Number of wounded 4T1 cell edges analyzed: 0 Pa (n = 28), 600 Pa (n = 33), 1,200 Pa (n = 24).

In line with recently published work which showed that the perturbation of intercellular adhesion (by inactivation of the E-cadherin gene) increased traction forces (Balasubramaniam *et al*., 2021), we found that 4T1 cell islands with low intercellular adhesion exert high traction stresses. Consistent with this, MCF10A islands which express high intercellular adhesion exert low traction forces. Compressive stress reduced traction forces for both cell types (**Fig. 5C**). When compressed MCF10A cells became jammed, the magnitude of traction stresses exerted by the cells decreased. Treating the cell islands with 1,200 Pa compressive load for 3 hours reduced traction stresses by 33.6% and strain energy by 52.1% in MCF10A cells (**Fig. 5D and 5E**). However, traction forces were also attenuated in compressed, unjammed 4T1 cells. Compressive stress largely obliterated cell traction forces in 4T1 islands as traction and strain energy decreased by 83.5% and 87.1%, respectively, compared to control islands (**Fig. 5D and 5E**). Compressive stress significantly elevated cell-cell adhesion in 4T1 cell islands (**Supplementary Figure 2**) while reducing cell-substrate stresses. Since microcontact printed islands lack the leading cell edges of coordinated migration, we can conclude that the traction forces of cells in the bulk of the monolayer were reduced by compressive stress regardless of the cell type we looked at or cell motility.

Our data from using micropatterned substrates indicated that cell-substrate stresses may not be the principal determinant of compression-driven unjamming in breast cancer cell migration. However, considering that traction forces were measured on microcontact printed islands, which do not permit coordinated migration, we cannot completely rule out the contribution of cell-substrate contraction during unjamming. Previous studies of collective cell migration during wound healing suggest that the leading edge of a cell sheet presents more cell-substrate adhesion than follower cells (Tse *et al*., 2012), which enables coordinated migration. In a wound healing assay, traction forces were localized to the MCF10A leading edge and were substantially reduced by compressive stress (**Fig. 5F and 5G**). Applying 1,200 Pa mechanical compression for 3 hours reduced traction by 64.3% in MCF10As and increased traction by 27.3% in 4T1s (**Fig. 5F and 5G**). High traction was exerted by control 4T1 cells throughout the cell layer, and mechanical compression increased traction stresses at the leading edge. Interestingly, despite an overall slight increase in traction stress with increasing compressive pressure for the regions analyzed (**Fig. 5G**), in compressed 4T1 cell sheets, cells at the leading edge exerted higher traction forces (**Fig. 5F**). Strain energy was found to diminish for MCF10A cell sheets and increase for 4T1 with higher compressive stress (**Fig. 5H**). Given these results, our data point to a potential differential leader-follower cell-substrate traction force response to compressive stress (i.e., increased traction of cells at the leading edge and reduced traction of cells in the bulk of the cell sheet) to be important in the unjamming behavior of 4T1 cells under mechanical compression.

### Theoretical simulation using the SPV model suggests distinct paths of jamming—unjamming

Our observed experimental data shows that there are two distinct responses to long-term compression from the two cell types of interest. The initially jammed 4T1 cells become unjammed under compression, whereas the MCF10A cells behave the opposite way, in a cell adhesion dependent manner. Seeking a theoretical explanation for this distinction, we investigate the self-propelled Voronoi (SPV) model and try to map the observed cell’s condition to a phase diagram of two model parameters (1) cell motility *v*_0_ and (2) target cell shape index *p*_0_ (The model and parameters are elaborated in more detail in the **Method** section). In the SPV model, the effect of cell-cell adhesion is captured by the parameter target shape index *p*_0_ (Bi *et al*., 2015). In this theoretical model, the effects of increased cell-cell adhesion lead to a higher value of *p*_0_. From the data for relative level of E-cad expression for MCF10A cells (**Figure 3B**), which show an insignificant difference in relative E-cad expression between control MCF10A and compressed MCF10A cells, we expect to see in the phase diagram that the difference in *p*_0_ of controlled and compressed MCF10A cells is small. This expectation is observed in **Figure 6**, where *p*_0_ of the mapped control MCF10A cells is 3.744, while of the mapped MCF10A compressed cells is 3.746. On the contrary, **Figure 3D** shows a significant difference in relative E-cad expression between control and compressed 4T1 cells. 4T1 cells under long-term compression express much more E-cad, which suggests a drastic increase in *p*_0_ for 4T1 cells under the effect of compression. This feature is also observed in the phase diagram, where *p*_0_ of 4T1 cells increases from 3.404 to 3.894 in response to compression. This E-cad – *p*_0_ comparison solidifies our experimental – theoretical mapping. Using a jammed—unjammed boundary proposed in ref. (Bi *et al*., 2016), we are able to see two distinct transition paths taken by 4T1 and MCF10A cells. The initially jammed 4T1 cells, under the effect of compression, become unjammed and cross the phase boundary from the jammed side. In contrast, MCF10A cells go from unjammed to jammed under the effect of compression (**Figure 6**).

**Figure 6.**
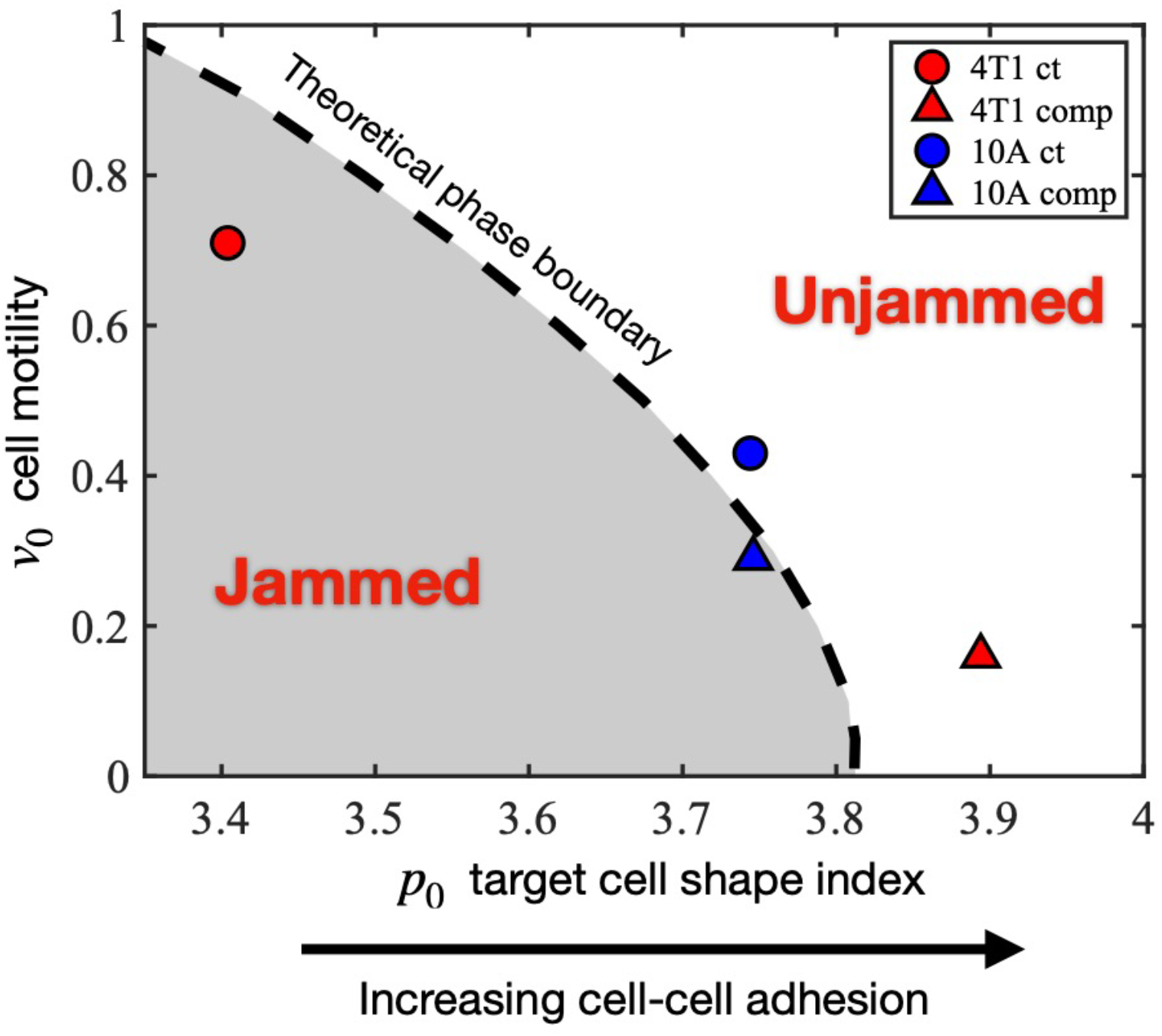
Mapping the experimentally observed tissue states to the theoretical jamming— unjamming phase diagram. The jamming—unjamming phase diagram is shown in terms of the two pertinent parameters of the SPV model: the single cell motility *v*_0_ and the target cell shape index *p*_0_. By mapping the experimentally observed cell traction forces and cell shapes to theoretical simulation results, the positions of 4T1 and 10A cells are placed on the phase diagram (see **Methods**).

## Discussion

In the present study, we show that cell-cell adhesion is the dominant mechanism for unjamming transitions in a dense, mechanically stressed monolayer. We first establish increased collective cell migration in a transition correlated with cell shape and distinct from EMT. Increased migratory behavior has traditionally been associated with EMT^31–33^, during which cells lose epithelial features and become mesenchymal (Nieto *et al*., 2016). However, in the transition we observe, cell motility increases, and the tissue fluidizes as the epithelial marker E-cadherin is upregulated and the mesenchymal marker vimentin is downregulated. These findings support a recently published characterization of the unjamming transition in which epithelial character is maintained without gaining mesenchymal character (Mitchel *et al*., 2020). As cell-cell adhesion increases, cells become fluid-like and rearrange more easily, increasing collective cell motion and accelerating wound repair. Despite 4T1 cells becoming more unjammed with compression, the cells exert lower traction forces. Attenuation of traction stresses in compressed MCF10A and 4T1 cells suggest that substrate traction may not be the dominant parameter in compression-induced unjamming. However, further investigation of traction forces during wound healing suggests that in a compressed cell sheet, leader cells exert increased traction forces, while follower cells present lower cell-substrate adhesion and higher cell-cell adhesion, allowing for greater structural rearrangements and fluid-like collective cell migration, where follower cells are more easily pulled by leading cells at the wound edge. Furthermore, we reveal that E-cadherin knockdown inhibits migratory behavior under mechanical compression, demonstrating that strong cell-cell adhesion, accompanied by increased levels of vinculin at cell-cell contacts, is important for regulating cellular unjamming. Our results suggest that cancer cells respond to compressive stress by becoming more elongated and strengthening cell-cell adhesions, leading to increased cohesion of the cell sheet, and enhanced collective migration.

One open question is how compressive stress leads to increased recruitment of E-cadherin to promote cell-cell adhesion. Mechanical stimuli are known to remodel epithelial cell-cell junctions by junction elongation and contraction mediated via mechanosensitive channels (Varadarajan *et al*., 2022). It is also well documented that adherens junctions become reinforced when cells are under tension (Borghi *et al*., 2012; Charras and Yap, 2018; Pinheiro and Bellaíche, 2018). We have previously found that mechanical compression activates mechanosensitive channel Piezo1 leading to calcium influx using a similar experimental setup (Luo *et al*., 2022). Interestingly, it was recently identified that Piezo1 directly binds to E-cadherin, and Piezo1 is tethered to the actin cytoskeleton via the cadherin-β-catenin complex (Wang *et al*., 2022). While these findings help support the idea of cell-cell junction stabilization during mechanical perturbation, it remains enigmatic how compressive stress could increase recruitment of E-cadherin to the cell membrane. Surface expression of E-cadherin would be a balance of endocytosis and vesicular trafficking to the plasma membrane (Kowalczyk and Nanes, 2012). Imbalance of trafficking of E-cadherin, for instance by reduced constitutive endocytosis, would alter the surface level of E-cadherin. We and others have shown that endocytosis is reduced when cell tension is elevated (Tan, Heureaux and Liu, 2015; Willy *et al*., 2017; Wu *et al*., 2017). Thus, it is plausible that elevated tension, due to the application of compressive stress, could slow down the turnover of E-cadherin on the cell surface. It becomes more interesting considering that cancer cells are softer than normal cells (Lee and Liu, 2015; Alibert, Goud and Manneville, 2017). A softer cell would become more deformed for a given compressive stress compared to a stiffer cell. In this way, the plasma membrane of cancer cells becomes more stretched by compressive stress, thereby resulting in reduced endocytosis and membrane protein turnover. Whether this could contribute to the increased E-cadherin level at cell-cell junctions in cancer cells in response to compressive stress remains to be further investigated.

It will also be of great interest to understand how significant upregulation of cell-cell adhesion translates to cells detaching from the solid tumor and migrating to adjacent tissues (Mendonsa, Na and Gumbiner, 2018). In cancer cells, heterogeneities in cell-cell adhesion within a tissue may be exacerbated by mechanical compression, allowing strongly adhesive cell clusters to move separately from their neighbors. Further studies in 3D cell spheroids will be necessary to decipher how these heterogeneities impact cell motility and thereby, migration and invasion. Furthermore, the sensitivity of different cell types to compressive stress may be related to characteristics of the nucleus. The nucleus is mechanosensitive, influences cellular force generation (Grosser *et al*., 2021), and may be actively involved in unjamming transitions. Interestingly, during wound healing, nuclear tension has been reported to decrease with distance from the wound edge, indicating higher tension in cells near the wound edge than in the bulk of the monolayer (Déjardin *et al*., 2020).

Our findings suggest a new physical picture of tumor development and cancer invasion, in which compressive stress inhibits the migratory ability of normal epithelial cells and allows cancer cells to migrate rapidly as a tightly connected group. The way cells react to mechanical compression is cell type-specific (Tse *et al*., 2012; Northcott *et al*., 2018; Luo *et al*., 2022) and drives the dense cancer cell monolayer to structurally rearrange in an unjamming transition that is not primarily driven by EMT. E-cadherin-dependent intercellular adhesion is the dominant regulator of cellular jamming and unjamming transitions driven by mechanical compression.

## Materials and Methods

### Cell culture

The human non-tumorigenic breast epithelial cell line MCF10A was a gift from Sofia Merajver (University of Michigan) and was obtained from Dr. Heppner at the Michigan Cancer Foundation where the cell line was originally developed. The mouse breast cancer cell line 4T1 was a gift from Lance Munn (Harvard Medical School) and was originally obtained from ATCC. The mouse breast cancer cell line 67NR was obtained from the Karmanos Cancer Institute (Detroit, MI). All cell lines were cultured in RPMI medium (Corning) supplemented with 10% FBS, except for MCF10A. MCF10A cells were cultured in DMEM/F12 medium (Corning) supplemented with 5% horse serum, 20 ng/ml epidermal growth factor (EGF), 0.5 μg/ml hydrocortisone, 100 ng/ml cholera toxin and 10 μg/ml insulin. Cells were cultured in a humidified atmosphere containing 5% CO_2_ at 37°C.

### Generation of E-cadherin knockdown 4T1 cells

4T1 cells expressing inducible shRNA knockdown of E-cadherin was generated using a transfer plasmid provided by Valerie Weaver at UCSF (Muncie *et al*., 2020). The transfer vector consisted of a modified pLKO.1 neo plasmid (Addgene) with expression of the shRNA sequences under control of 3× copies of the lac operator and a copy of the mNeonGreen fluorescence protein. Lentiviruses were generated by transfecting HEK 293T cells with the transfer vector, psPAX2 packaging vector, and pMD2.G envelope vector. Viral supernatant was collected 48 h after transfection. 4T1 cells were transduced in RPMI medium and after 24 h, shE-cadherin cells were selected with 200 μg/ml G-418 (Sigma). E-cadherin knockdown was induced in shE-cadherin cells by adding 200 μM isopropyl-β-D-thiogalactoside (IPTG; Sigma) 72 h prior to experiments.

### Mechanical compression using an in vitro compression setup

Vertical compression was applied by adding a weight of 26 or 52 g over an area of 426 mm^2^ to achieve stresses of 600 or 1,200 Pa, respectively, using a previously established setup (Tse *et al*., 2012; Luo *et al*., 2022). The fixed weight applies a constant stress to a UV-treated 1% agarose gel cushion in contact with cells. The agarose gel allows nutrient and oxygen diffusion. The agarose gel was used without the weight as a negative control. Samples for live imaging, immunofluorescence staining, and RT-qPCR were compressed for the specified durations under cell culture conditions.

### In vitro scratch-wound assay and time-lapse imaging

Cells were plated onto 6 well plates 72 h prior to experiments and grown to confluence. MCF10A and 4T1 cells were serum-starved in DMEM/F12 without horse serum and EGF and RPMI medium without FBS, respectively, for 4 h prior to experiments. Cells were then incubated with Hoechst 33342 diluted in PBS for 30 min at 37°C. Cells were washed with PBS and placed in complete medium. Scratches were created using a p-200 pipette tip to induce migration. After wounding, vertical compressive stresses of 0, 600 and 1,200 Pa were applied.

Images of the wound area were captured using a 4× objective at 30 min time points for 15 h using fluorescence microscopy on a Cytation 5 automated plate reader. Collective cell migration was quantified by measuring the area between wound edges using MATLAB. Spatiotemporal velocity maps were generated using the software AveMap+ (Deforet *et al*., 2012). Results were collected from 3 independent experiments.

### Immunofluorescence staining

Cells on glass coverslips were washed with PBS and fixed with 4% paraformaldehyde for 10 minutes, washed with PBS, and permeabilized with 0.1% Triton X-100 in PBS for 10 minutes. Cells were washed with PBS and blocked with 3% BSA in PBS for 1 h. Cells were incubated with a rabbit anti-E-cadherin antibody at 1:400 (Cell Signaling) and a primary mouse anti-vinculin antibody at 1:800 (Sigma-Aldrich) in 3% BSA overnight at 4°C. Cells were washed 3× with PBS and incubated with DAPI, phalloidin, and secondary antibodies in 3% BSA for 1 h at room temperature. Cells were washed 3× with PBS, mounted onto glass slides with Fluoromount-G (Invitrogen), and imaged by spinning disk confocal microscopy. Fluorescence levels relative to the control condition were quantified. E-cadherin at cell–cell contacts was quantified by measuring fluorescence intensity at the cell membrane.

### Confocal microscopy

Images of immunostained cells were taken using an oil immersion Plan-Apochromat 60 x/1.4 NA objective on an inverted microscope (Olympus IX-81) equipped with an iXON3 EMCCD camera (Andor Technology), AOTF-controlled lasers (Andor Technology), and a Yokogawa CSU-X1 spinning disk confocal. Acquisition of images was controlled by MetaMorph (Molecular Devices). Single and z-stack images of cells fluorescently labeled for DAPI, F-actin (by 488-phalloidin), E-cadherin, and vinculin were captured with 405 nm, 488 nm, 561 nm, and 640 nm excitations, respectively, at exposure times of 200-500 ms.

### Quantification of cell and nuclear shapes

Immunofluorescence images were acquired as described above. For each pressure condition, more than 200 cells and nuclei were manually traced using ImageJ software (National Institutes of Health) from ten different fields of view. The cell and nuclear shape indices were computed for each traced cell and nucleus, respectively.

### RNA extraction and RT-qPCR

RNA was extracted using the RNeasy micro kit (Qiagen). RNA quality and quantity were measured using a NanoDrop 1000 spectrophotometer. Reverse transcription was performed using the iScript cDNA synthesis kit (Bio-Rad). qPCR assays were conducted using SYBR Green (Bio-Rad) and specific primers quantifying *gapdh, cdh1, cdh2*, and *vim* (OriGene) on the Bio-Rad CFX thermocycler. *Gapdh* was used as a control for quantifying relative gene expression. Mean C_t_ values from duplicates were used to calculate ΔC_t_ values normalized to GAPDH. Relative transcript levels were determined by calculating the change between ΔC_t_ values of control and compressed samples as ΔΔC_t_ and calculating 2^-ΔΔC^t. Results were collected in duplicates from 3 independent experiments.

### Fabrication of hybrid silicone substrates

CY52-276 A/B (Dow Corning) with an A:B ratio of 1:1 was cast in 35 mm glass bottom dishes. After 10 min of degassing, the soft silicone substrates were cured on a hot plate at 70°C for 30 min. The substrates were then exposed to deep UV light for 5 min. 19 mg EDC (1-Ethyl-3-(3-dimethylaminopropyl)-carbodiimide) (Thermo Fisher), 11 mg sulfo-NHS (N-Hydroxysulfosuccinimide) (Thermo Fisher), and 15 μl 2% w/v 0.5 μm carboxylate fluorescent beads (Thermo Fisher) were added to 1 ml DI water. The substrates were incubated with this suspension for 30 min to conjugate fluorescent beads to the surface of the soft silicone (Bashirzadeh *et al*., 2018; Bashirzadeh, Qian and Maruthamuthu, 2018).

### Micropatterning of silicone substrates

Micropatterned substrates were made with standard soft lithography technique to create a silicon master mold. Polydimethylsiloxane (PDMS) was prepared by mixing Sylgard-184 elastomer and curing agent (Dow Corning) in a 10:1 (w/w) ratio. After 10 min of degassing, the PDMS mixture was poured over the master and cured overnight at 60°C. PDMS stamps were incubated with collagen I from rat tail (Corning) at a concentration of 50 μg/ml for 1 h. Soft silicone substrates were UV-treated for 5 min and then immediately placed in contact with the stamps. Printed substrates were passivated with anti-adherence rinsing solution (STEMCELL Technologies) for 1 h.

### Traction force microscopy

Cells were seeded on printed soft silicone substrates 48 hours prior to experiments. For control and compressed samples, cell islands labeled by Hoechst 33342 and red fluorescence beads on the substrate surface were imaged before and after removal of cells using 10% sodium dodecyl sulfate (SDS). PIVlab (Thielicke and Stamhuis, 2014) was used to process image pairs (bead images before and after cell removal) and then used with the fast Fourier transform window deformation method to quantify the displacement of the beads, resulting in a displacement vector field. The Young’s modulus of the substrate was previously measured to be 7.2 kPa using sphere indentation (Bashirzadeh *et al*., 2019). Fourier transform traction cytometry was used to compute the traction stress field using MATLAB (Butler *et al*., 2002; Schwarz *et al*., 2002; Sabass *et al*., 2008; Plotnikov *et al*., 2014).

### Statistical analysis

Statistical analysis was carried out in Origin and performed with one-way ANOVA followed by Tukey post-hoc multiple comparisons test. Results were collected from 3 independent experiments and plotted as mean ± S.D. or shown as boxplots. Statistical significance was denoted by asterisks in the figure panels, with ∗= *p* < 0.05, ∗∗= *p* < 0.01, ∗∗∗= *p* < 0.001.

### Theoretical model and simulation details

Cells in a 2D monolayer in the SPV model (Bi *et al*., 2016) are represented by polygons determined from a Voronoi tessellation of their center positions (*r*_*i*_). The center of each cell obeys the over-damped equation of motion

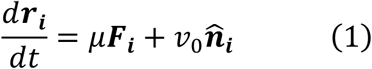

The first term on the right hand side (RHS) of Eq. (1) comes from cell-cell interactions and the mechanical behavior of a single cell. Here *µ* is the single cell mobility constant and the interaction force is given by *F*_*i*_ = −∇_*i*_*E*, where *E* is the total mechanical energy of the tissue given by

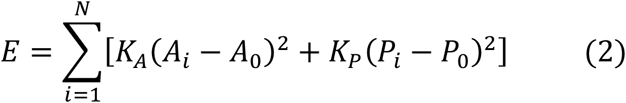

In this equation, *A*_*i*_ and *P*_*i*_ are area and perimeter of cell *i*th, *A*_0_ and *P*_0_ are the cell preferred area and perimeter, respectively. *K*_*A*_ and *K*_*p*_ are the area and perimeter moduli. The term involving cell area models cell’s incompressibility and monolayer’s resistance to height fluctuation. The quadratic term in perimeter results from active contractility of subcellular cortex. The linear term in perimeter is a combination of cortical tension and membrane tension due to cell-cell adhesion. The membrane line tension can be reduced by either increasing cell-cell adhesion, which encourages the cell to lengthen its shared edges with its neighbors, or by reducing actin-myosin contractility. Therefore, *P*_0_ is positively correlated to cell-cell adhesion and negatively correlated to cell contractility (Farhadifar *et al*., 2007). For simplicity, we assume that the contribution of cell-cell adhesion to line tension is greater than the contribution of actin-myosin contractility, so *P*_0_ is positive as a consequence. N is the total number of cells in the monolayer. To nondimensionalize cell shape quantities, we adapt a target shape index parameter 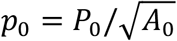.

In addition to the interaction force, cells are driven by a self-propelled force due to their own polarized motility. In the SPV model, this is captured by the second term on the RHS of Eq. (1), where *v*_0_/*µ* is the self-propulsive force magnitude. For each cell, this force acts along a polarization vector, given by

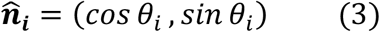

that determines the direction of the self-propelled force. The polarity of cells is stochastic and obeys rotational Brownian dynamics, given by

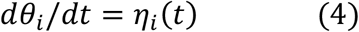

where η_*i*_(*t*) is a white noise process with zero mean and variance 2*D*_*r*_, given by

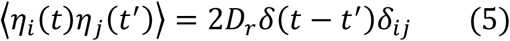

The magnitude of the rotational noise *D*_*r*_ defines a time scale *Ä* = 1/*D*_*r*_ of persistent motion.

We employ an open source implementation of the SPV model to perform CellGPU (Sussman, 2017) simulations of 900 cells for 5 million time steps with *dt* = 0.05 at each parameter of *p*_0_, *v*_0_ shown in **Fig. 6**. We choose *D*_*r*_ = 1, *K*_*A*_ = 1, *K*_*p*_ = 1, *A*_0_ = 1.

In the SPV model, the traction force or the total force exerted by the cell onto the substrate is given by a sum of the viscous friction between the cell and the substrate and the self-propulsive force

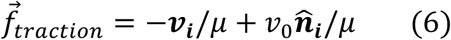

Since the net force on each cell is balanced according to Equation (1), the traction 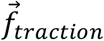 is always equivalent to the interaction force. Therefore, we use the interaction force to determine the magnitude of the traction of every cell in the system.

### Mapping condition of cells in vitro to theoretical simulation

To connect the experimental results to our theoretical model, we use (1) the observed cell aspect ratio (AR) values and (2) the cell traction forces to map the conditions of 4T1 and MCF10A cells to theoretical simulation parameters.

We performed simulations of the SPV model for a large range of *v*_0_- *p*_0_ values. For each simulation step, we compute the cell AR of each cell using the open-source function. The mean aspect ratio for each simulation was then calculated by averaging the AR over every cell over simulation time steps. A contour map of mean AR in the *v*_0_-*p*_0_ space is shown in (Supplementary **Fig. 4**). On the other hand, the experimental values of mean AR for each condition are labeled in the *v*_0_-*p*_0_ space and the locus of (*v*_0_, *p*_0_) pairs having these values of AR form contours. However, this alone does not give a definitive value of (*v*_0_, *p*_0_). The jamming phase boundary shown in **Fig. 6** is adapted from ref. (Bi *et al*., 2016) and demarcates solid-like states from fluid-like states, which is determined from the collective diffusive property at the tissue level. We next map the experimental values of cell tractions to the theoretical ones.

The average traction magnitude of the simulation systems that are on these contours is computed. We denoted the average traction magnitude to be T. To connect experimental traction values to simulations, we choose the traction values of 4T1 cells in the compressed case to map to the simulation traction value at (*p*_0_ = 3.404, *v*_0_ = 0.71). As a result, T(4T1 control) = 4.4790 was mapped to simulation traction of T(*p*_0_ = 3.404, *v*_0_ = 0.71) = 0.626. This is an assumption in the analysis. However, the particular choice of this mapping does not influence the final qualitative conclusion in our work. The simulation traction magnitude of the system at this position is divided by the experimentally measured 4T1 traction to obtain the conversion factor. The traction values of other cell types and conditions are converted to simulation units by multiplying them by the factor of conversion. The positions of other cell conditions on the phase diagram are then determined by matching both the AR and traction values between the experiment and simulation.

## Acknowledgements

The authors thank Jin-Ah Park and Jeffrey Fredberg of Harvard University for helpful discussions. The authors thank Lance Munn of Harvard Medical School for providing 4T1 cells. A.P.L. acknowledges support from the National Institutes of Health (grant no. R21GM134167) and the National Science Foundation (grant no. CMMI-1927803). D.B. and A.N. acknowledge support from the National Science Foundation (grant no. DMR-2046683), the Alfred P. Sloan Foundation, MathWorks Microgrants, and the Northeastern University Discovery Cluster. G.C. acknowledges support from the University of Michigan Rackham Merit Fellowship.

