## Supplemental Information for "Compressive stress drives adhesion-dependent unjamming transitions in breast cancer cell migration"

### *Supplementary Material*

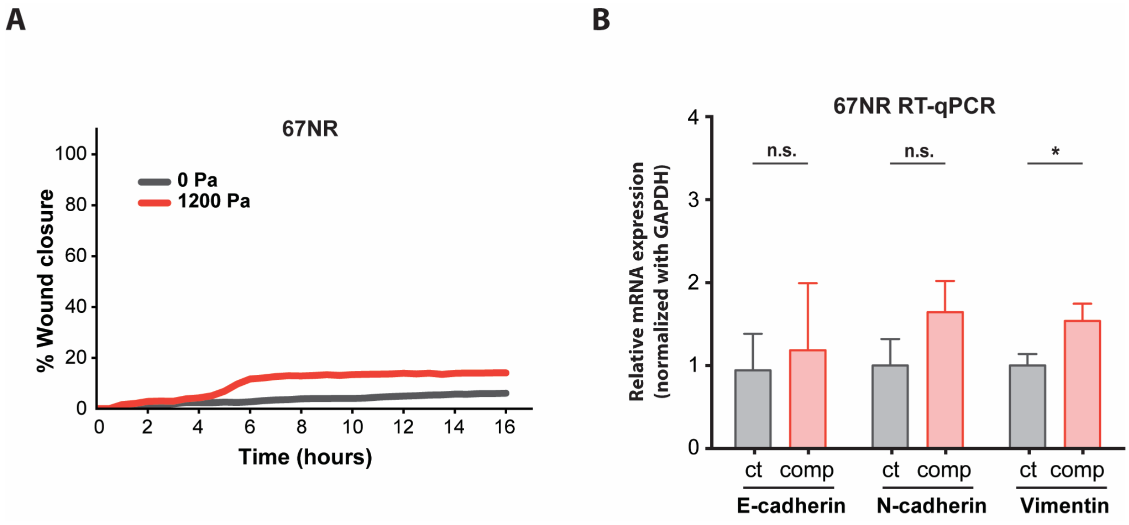


**Supplementary Figure 1. (A)** Quantification of wound area for 67NR cells with and without compression. Mean wound area at each time point is plotted from 3 independent replicates as a representative trace. **(B)** qPCR analysis of E-cadherin, N-cadherin, and vimentin mRNA levels with and without compression (1,200 Pa). Transcript levels are calculated using the $\Delta\Delta C_t$ method normalized to GAPDH. Mean mRNA level ± S.D. is plotted from 3 independent experiments with duplicates per experiment.

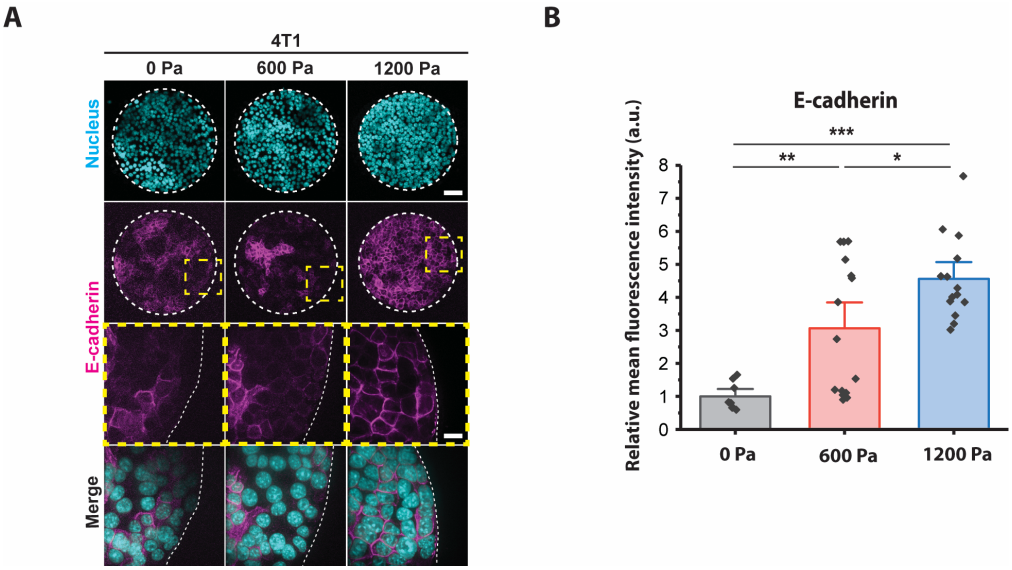


**Supplementary Figure 2.** **(A)** Representative microscopy images of 4T1 cell islands labeled with DAPI and an E-cadherin antibody. Micropatterned cell islands are exposed to specified stresses for 12 h. Scale bars, 80 µm (top) and 20 µm (bottom). **(B)** Quantification of relative E-cadherin fluorescence at 67NR cell-cell contacts. Mean fluorescence intensity at the cell membrane ± S.D. is plotted from 3 independent replicates (n = 8-15).

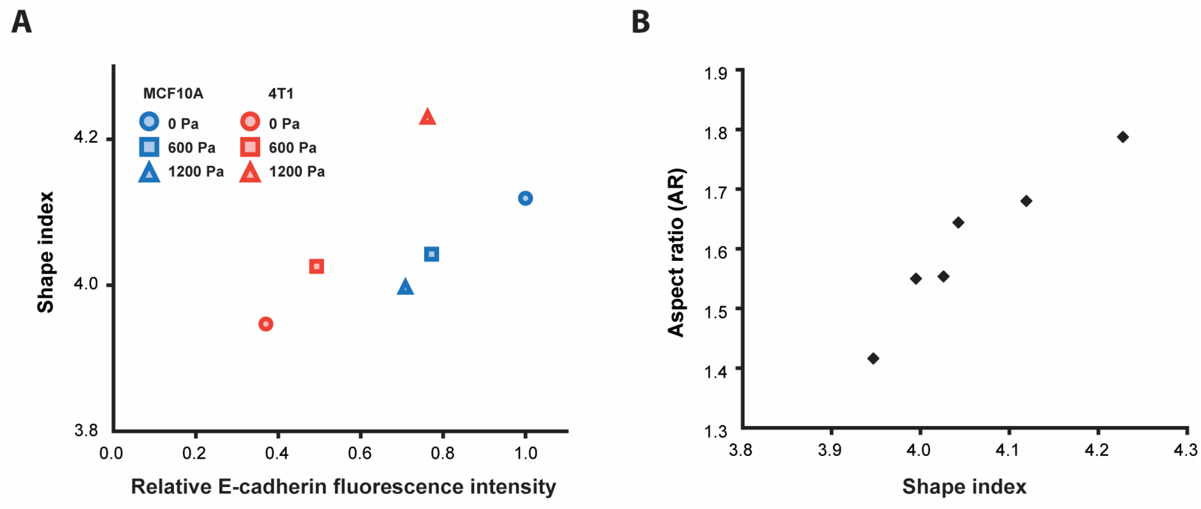


**Supplementary Figure 3. (A)** Plot of E-cadherin fluorescence level vs. cell shape index for MCF10A and 4T1 cells under specified compressive stresses. As the relative E-cadherin fluorescence intensity at cell-cell contacts is elevated, shape index increases as well. **(B)** Plot of shape index, which emphasizes perimeter, vs. aspect ratio (AR), which emphasizes elongation, shows that substantial increases in AR are accompanied by smaller increases in shape index, resulting in elongated cells with straight edges.

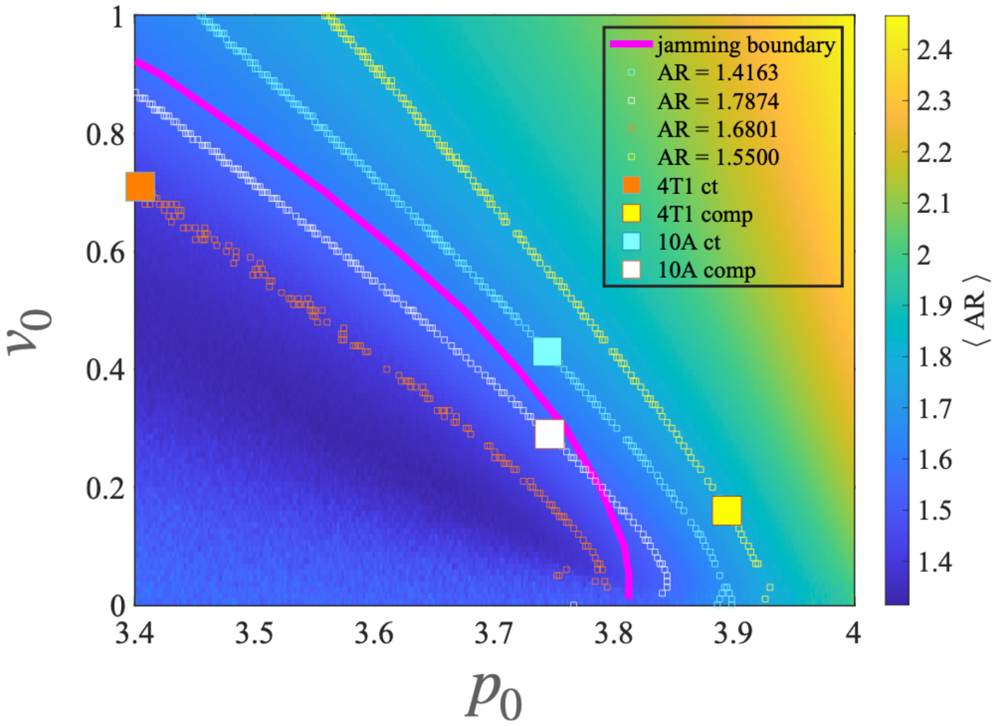


**Supplementary Figure 4.** Contour plot of mean AR at different $v_0$ and $p_0$ is represented as a color plot. The empty squares represent contour lines with the particular value of AR observed in experiments. The filled squares represent different cell lines and experimental conditions mapped by extrapolating from cell traction.

**Description of Additional Supplementary Files**

Supplementary Movie 1. **Collective cell migration of control MCF10A cells in a scratch wound assay.** A confluent monolayer of MCF10A is scratched and the wound edge is tracked by time lapse fluorescence imaging. Cell nuclei are labeled with Hoechst 33342.

Supplementary Movie 2. **Collective cell migration of compressed MCF10A cells in a scratch wound assay.** A confluent monolayer of MCF10A is scratched and compressed (1,200 Pa). The wound edge is tracked by time lapse fluorescence imaging. Cell nuclei are labeled with Hoechst 33342.

Supplementary Movie 3. **Collective cell migration of control 4T1 cells in a scratch wound assay.** A confluent monolayer of 4T1 is scratched and the wound edge is tracked by time lapse fluorescence imaging. Cell nuclei are labeled with Hoechst 33342.

Supplementary Movie 4. **Collective cell migration of compressed 4T1 cells in a scratch wound assay.** A confluent monolayer of 4T1 is scratched and compressed (1,200 Pa). The wound edge is tracked by time lapse fluorescence imaging. Cell nuclei are labeled with Hoechst 33342.
